# Curcumin Inhibits Zika Virus NS3 Helicase Activity

**DOI:** 10.1101/2024.01.03.573903

**Authors:** Ankur Kumar, Aparna Bhardwaj, Richa Joshi, Taniya Bhardwaj, Rajanish Giri

## Abstract

In the last decades, many efforts have been put forward in search of inhibitors/drugs specific to ZIKV-infected patients. These inhibitors include approved drugs, natural products, nucleosides, small molecules, and many others to inhibit Zika virus replication in cell and mouse models. In this regard, natural product curcumin has been widely explored in various disease conditions and found to have inhibition potential against ZIKV replication as well. Further, we investigated its inhibitory ability against the ZIKV NS3 helicase. We found that the curcumin interacts with the NS3 helicase domain by forming stable contacts at NTPase and RNA binding sites during molecular dynamic simulation. Furthermore, curcumin inhibits NTPase activity of NS3 helicase with an IC_50_ of 64.42 μM. Altogether, the mechanistic insight into the action of curcumin will enlighten further drug discovery research in finding novel molecules.

## 1. INTRODUCTION

Flaviviral NS3 helicase is an indispensable protein for unwinding viral RNA during replication. It uses ATP as a substrate for NTPase activity, providing energy for the helicase for viral genome synthesis in coordination with NS5 polymerase [1,2]. The ZIKV NS3 helicase is observed to be a monomer in a solution having three subdomains: DI (residues 175-332), DII (residues 333-481), DIII (residues 482-617) [3,4]. An ATP/Mg^2+^ binding site is located in between domain DI and DII, where residues Gly197, Lys200, Thr201, Arg202, Glu286, Asn330, Arg459, and Arg462 are involved in the binding interaction. It has been shown that residues 193-202 and residues 249-255 have a high B-factor, making these regions significantly flexible [4]. Further, eight structural motifs on the ZIKV NS3 helicase domain are involved in ATP hydrolysis, RNA binding, and establishing a connection between the binding sites [2,4,5]. Due to the conserved catalytic residues, the substrate-, RNA-, and ATP/Mg^2+^-binding sites act as important targets for the inhibitor design [3] [4].

Viral helicases have been an important target for structure-based drug discovery to find antiviral compounds. For example, some small molecules and nucleoside analogs have been identified previously to inhibit the helicase activity of HCV [6–8]. Similarly, small molecules that contain benzothiazole and pyrrolone scaffolds have been reported to inhibit the NTPase activity of NS3 helicase in DENV [9]. The helicase activity of flaviviral helicases from YFV, DENV, and WNV is shown to be inhibited by ivermectin [10]. A natural compound, epigallocatechin-3-gallate, inhibited ZIKV NS3 helicase activity at a low micromolar range [11]. Curcumin is a naturally occurring lipophilic polyphenolic compound extracted from turmeric roots that acts as a potent antioxidant, anti-inflammatory, anti-viral, and anti-bacterial compound [12,13]. Moreover, it was also identified as an inhibitor of a methicillin-resistant strain of *Staphylococcus aureus* [14]. Besides these activities, curcumin also exhibits anti-neurodegenerative and anti-cancer properties and affects cellular senescence [15]. Curcumin exerts its anti-viral effects through several means, such as obstructing viral attachment, entry into host cells, replication, and degradation of viral proteins [16]. Among flaviviruses, it has been demonstrated to inhibit DENV entry and replication, and, in addition, it also impedes the NS3 protease function [17].

Further, in this study, we focused on investigating the molecular interaction of curcumin with ZIKV NS3 helicase. The previously identified NTPase and RNA binding sites [18] were used here as druggable sites to dock the curcumin. In our *in silico* studies, the curcumin formed stable interactions at NTPase and RNA binding sites. Further, the curcumin inhibited the NTPase activity of NS3 helicase. Thus, the molecular interaction between curcumin and NS3 helicase will enlighten drug discovery research in finding novel molecules for the Zika virus and other closely related flaviviruses.

## 2. MATERIAL AND METHOD

### 2.1. Software

Schrodinger software package was used to perform all the computational studies in this article.

### 2.2. Receptor grid generation

We used the previously determined X-ray crystal structure of NS3 helicase (PDB ID: 5GJC and 5GJB) for our in-silico studies [18]. At first, the protein structure was prepared using the protein preparation wizard in the Glide module of Maestro 11.2 version, where the protein structure was preprocessed and optimized at pH 7, as described in our previous article [11]. Then, the optimized structure was minimized using OPLS 2005 forcefield by converging heavy atoms to RMSD of 0.30 Å. Further, the minimized structure was used to create a grid on the NTPase site and at the RNA binding site by employing receptor grid generation of the Glide module. In the crystal structure 5GJC, the centroid of the bound ATP at the NTPase site was chosen to generate a grid with a size length of 20 Å. Similarly, in 5GJB, the centroid of the bound RNA at the RNA binding site was used to generate a grid over RNA binding site with a size length of 20 Å. We have taken a scaling factor of 1 and a partial charge cutoff of 0.25 to scale the van der Waals radius of the receptor. We also prepared a grid on the NS2B-NS3 protease (5LC0) active site using the same parameter by choosing an enclosing box at the centroid of the catalytic triad by selecting His51, Asp75, and Ser135 residues. In the case of MTase and RdRp, we utilized the same receptor grid described in our previously published article [19].

### 2.3. Ligand preparation

We took SDF file of the curcumin from the PubChem database and further prepared it in LigPrep module in the Mastero 11.2 version. We used OPLS_2005 force field to prepare the ligand, and the preparation was performed by generating a possible ionization state at pH 7 using Epik. The stereoisomers were generated by retaining specified chirality in the molecule. Further, we chose to desalt and generate tautomers for curcumin.

### 2.4. Molecular docking

We use the Glide module of the Maestro 11.2 version to perform the ligand docking. The prepared structure of curcumin was docked to the prepared receptor grid (described in section 2.2) at NTPase and RNA binding site of NS3 helicase, the active site of NS2B-NS3 protease, the druggable site of MTase, and the druggable site of RdRp. We used a scaling factor of 0.80 and a partial charge cutoff of 0.15 to scale the van der Waals radii of ligand atoms. We chose an extra precision (XP) docking protocol while selecting a flexible ligand sampling method (added sample nitrogen inversion and sample ring conformation). Further, we used bias sampling of torsions for amides to penalize nonpolar conformation. A 2.5 kcal/mol of energy window was set for ring sampling. We kept a distance-dependent dielectric constant of 2 for energy minimization with 100 minimization steps and used OPLS_2005 forcefield for docking protocol. We perform a post-docking minimization step to include five poses per ligand.

### 2.5. Binding-free energy calculation

We use a similar approach to calculate the binding-free energy calculation of the protein-ligand complex as described in our previous article [20–22]. We use the XP data file from the docking result as the input to obtain binding free energy for the curcumin-protein complex. The obtained poses after XP docking were subjected to the Prime module of Maestro 11.2, which uses MM-GBSA method to calculate the binding energy [23]. We utilized VSGB solvation model and OPLS_2005 force field to run the MM-GBSA.

### 2.6. Molecular dynamics simulation

The stability of the interaction between curcumin and NS3 helicase (NTPase site and RNA binding site) was checked further by subjecting the curcumin-NS3 helicase complex to Molecular dynamics (MD) simulation for 200 ns. The Desmond simulation package was used for MD simulation by following previously described protocols [21]. In the system builder, the curcumin-NS3 helicase complex was enclosed in the center of an orthorhombic box with an edge distance of 10 L, and this box was filled with TIP4P predefined solvent model. The sodium ion was recalculated for the system, and 0.15 M NaCl salt was added to neutralize uniform electrostatic distribution. The system was further minimized using 5000 iterations (1 kcal/mol/L convergence threshold) containing the steepest descent method of 10 steps and three LBFGS vectors until a gradient threshold is reached up to 25 kcal/mol/L. The cutoff coulombic radius of interaction was set up to 9 L. Finally, the simulation run was carried out for 200 ns with NPT ensemble class having 300 K temperature and 1.01 bar pressure. We chose to relax the model system before simulation. Nose-Hoover thermostat (relaxation time: 1ps, number of groups:1) and Martyna-Tobias-Klein (MTK) barostat methods (relaxation time: 2ps and isotropic coupling style) were used for temperature and pressure coupling throughout the simulations. The cutoff coulombic radius of interaction was set up to 9 L. When the simulation was completed, the simulation parameter was analyzed using by simulation event analysis run and a simulation interaction diagram analysis run.

### 2.7. NS3 helicase inhibition assay

NS3 helicase activity and inhibition assay (see the detailed protocol of protein purification, activity, and inhibition assay as described in a previous report by Kumar et al., 2020 [11]) were performed in Tris-HCl buffer (pH 7.4). For the NTPase inhibition assay, a required amount of NS3 helicase purified protein was incubated for 15 minutes with various concentrations of curcumin (purchased from Sigma-Aldrich; 50 mM stock of curcumin was prepared in 100% DMSO). Afterward, the curcumin-NS3 helicase complex (20 μl) was mixed with the substrate (20 μl), ATP, and incubated for 20 minutes. Then, malachite green reagent (160 μl) was added to terminate the reaction (see malachite green reagent preparation in previous report by Kumar et al., 2020 [11]). After 5 minutes, absorbance was monitored at 630 nm using a multi-well plate reader (infiniteM200PRO: TECAN). The final NS3 helicase concentration in the reaction was kept at 80 nM. The reaction was performed in triplicates on a 96-well transparent plate. For the well having no curcumin, we use DMSO, which corresponds to the same DMSO percentage for the well having higher Curcumin concentration in the respective set of experiments. Absorbance was converted to the release of phosphate using a standard curve plotted using free phosphate. Further, the phosphate release per minute (V) was plotted as the function of substrate [µM] to compute kinetic parameters and IC_50_ using GraphPad Prism.

## 3. RESULT

### 3.1. Molecular Docking of curcumin at NTPase site of NS3 helicase

During viral RNA synthesis, the NTPase site acts as the binding site of ATP, and after hydrolysis, it provides energy to RNA unwinding process. Therefore, blocking this site will lead to the inhibition of NS3 helicase function. First, we see the molecular interaction of curcumin at the NTPase site of the NS3 helicase. We used the 5GJC crystal structure of NS3 helicase, where a grid was prepared on the centroid of the ATP binding site (Figure 1A). Further, curcumin was docked on the receptor grid by employing an extra precision (XP) docking protocol. Curcumin shows a docking score of −4.93 kcal/mol and stabilizes itself at the NTPase site via H-bond (Figure 1B). This H-bond is formed between the hydroxyl group of Curcumin and the Phe418 residue of NS3 helicase. Besides H-bonds, the curcumin stabilizes at this site with the hydrophobic interactions via Pro196, Ala198, Ala420, and Pro464 residues (Figure 1C). Still, the molecular interaction of curcumin with NS3 helicase needs further experimental validation by NMR, and the crystal structure of the protein-ligand complex.

**Figure 1:**
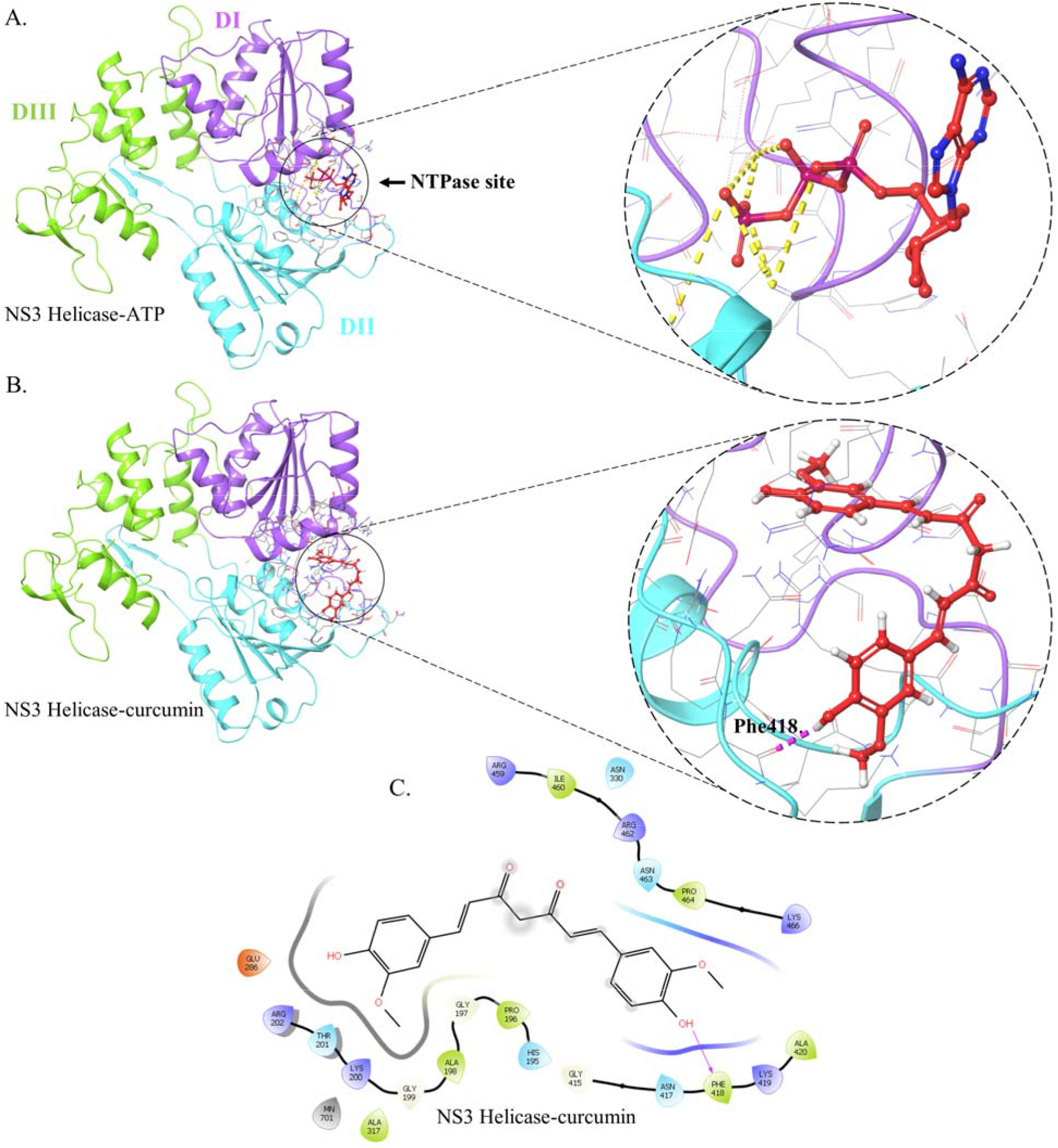
Molecular interaction of curcumin with the NTPase site of NS3 helicase in docking. (A.) 5GJC crystal structure of NS3 helicase representing the bound ATP (shown in red) at the NTPase site. Three domains, DI (178-332), DII (333-481), and DII (482-617) of NS3 helicase, are represented in magenta, cyan, and green color, respectively. At the NTPase site, the phosphate group oxygen of ATP forms a salt bridge (yellow dashed line) with Lys200, Arg459, and Arg462 residues. Additionally, phosphate group oxygen also forms salt bridges with Mn^2+^. (B) The three-dimensional interaction pose of curcumin and NS3 helicase (5GJC) represents curcumin’s interaction with the amino acid residues at the NTPase site. The pink dashed line represents the interaction of curcumin via H-bond with Phe418 of NS3 helicase. (C) Two-dimensional pose of curcumin at the NTPase site of NS3 helicase representing its interaction with the amino acid residues. The hydrophobic residues of helicase are represented in green color. The pink arrow indicates the H-bond between the curcumin and the amino acid residue of NS3 helicase (5GJC).

### 3.2. Curcumin forms stable interaction at NTPase site of NS3 helicase during MD simulation

In the docking, the curcumin showed interaction at NTPase site of NS3 helicase via H-bond and hydrophobic interaction. So, further, we checked the binding stability of curcumin and the conformational flexibility of NS3 helicase (5GJC) upon binding of Curcumin to its NTPase site by subjecting ligand-protein complex to simulation for 200 ns (Figure 2). During the simulation, the C-α RMSD of curcumin-bound NS3 helicase was observed between 1.29-2.51 Å (Figure 2A), which is a little higher than the unbound NS3 helicase (1.17-2.37 Å). The average RMSD of NS3 helicase is 1.92 Å or 1.99 Å in the absence or presence of curcumin, respectively. These findings suggest that the NS3 helicase exhibited a stable conformation when curcumin binds at the NTPase site and did not show significant variation compared to unbound NS3 helicase. Furthermore, the ordered regions (helix or strand) of NS3 helicase showed lower RMSF than the disorder region (coil or loop), either in the presence or absence of curcumin, which further complements the binding of EGCG at the NTPase site, as reported in our previous literature [11]. Notably, when curcumin binds at the NTPase site, the regions between 244-256, 271-279, 412-419, 434-439, and 570-580 showed low RMSF values compared to unbound NS3 helicase. Val250, Arg275, Gly415, Gly437, and Glu573 residues corresponding to the respective five regions in the NS3 helicase showed higher RMSF without curcumin (Figure 2B).

**Figure 2:**
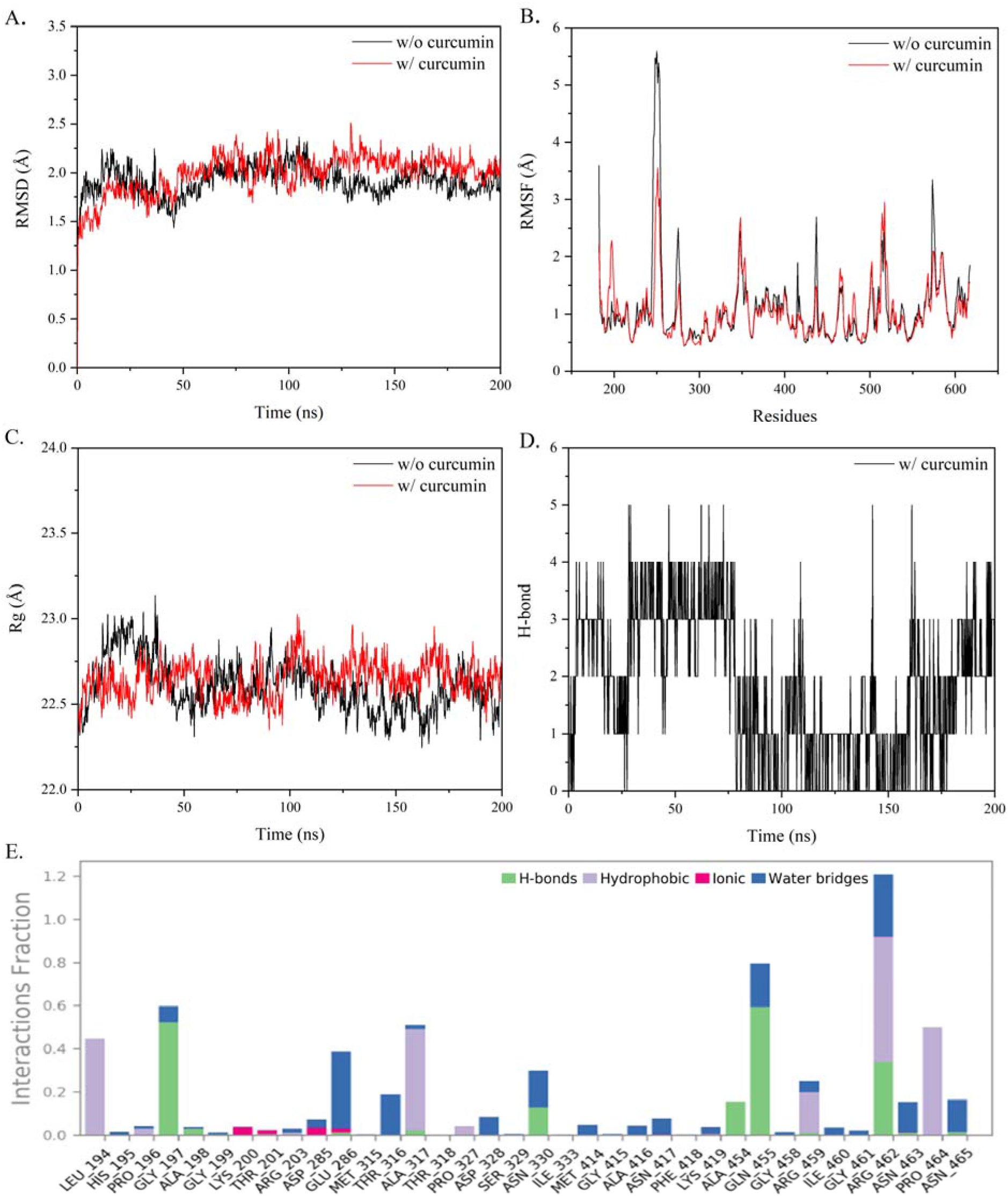
MD simulation of NS3 helicase when curcumin interacted with NTPase site. (A) The RMSD of NS3 helicase in the presence (red colored line) and the absence (black colored line) of curcumin during simulation. (B) The RMSF value of each residue of NS3 helicase in the presence (red colored line) and the absence (black colored line) of the curcumin. (C) The Rg of NS3 helicase in the presence (red colored line) and the absence (black colored line) of curcumin. (D) The number of hydrogen bonds between curcumin and the NTPase site of NS3 helicase. (E) The fraction of interaction of NTPase site of NS3 helicase residues with the curcumin by H-bond, hydrophobic interaction, ionic bond, and water bridges during simulation of 200 ns. A fraction of 1 represents that the particular residue of helicase contacted with curcumin for 100 % of the simulation time. [5GJC was used to dock the curcumin at NTPase site of NS3 helicase]

The Rg value of NS3 helicase bound with curcumin shows a slightly variable trend compared to unbound helicase (Figure 2C). Still, the average Rg value of NS3 helicase is nearly similar, either in the presence (22.65 Å) or absence of curcumin (22.60 Å). NS3 helicase in the absence of curcumin showed the total secondary structure element to be 38.31 %, where helix and strand are 24.63 % and 13.68 %, respectively (Figure S1). When bound with Curcumin, there is a little increase in the total secondary structure element (Figure S2) of NS3 helicase (Total SSE 39.05 %; Helix 24.66 % and Strand 14.39 %). Although curcumin forms only one H-bond with Phe418 in the docking, it interacted with many residues during the simulation. It forms a maximum of five H-bonds at a time in a few of the trajectory’s frames (Figure 2D). The curcumin interacted with 36 residues and maintained contact with variable fractions of simulation time of 200 ns (Figure 2E & S3). Gly197, Ala198, Gly199, Thr201, Arg203, Glu286, Ala317, Asn330, Phe418, Lys419, Ala454, Gln455, Arg459, Arg462, Asn463, and Asn465 are involved in forming H-bond contacts with the curcumin. Lys200, Thr201, Asp285, Glu286, Asn417, Lys419, Arg462, and Asn463 are involved in forming ionic bonds with curcumin. Leu194, Pro196, Arg203, Ala317, Pro327, Ile333, Arg459, Arg462, and Pro464 are involved in forming hydrophobic bonds with curcumin. His195, Pro196, Gly197, Ala198, Gly199, Arg203, Asp285, Glu286, Met315, Thr316, Ala317, Asp328, Ser329, Asn330, Met414, Gly415, Ala416, Asn417, Lys419, Gln455, Gly458, Arg459, Ile460, Gly461, Arg462, Asn463, and Asn465 are involved in forming water bridges with curcumin. Among all the contacts between curcumin and NTPase site of NS3 helicase, residues Leu194, Gly197, Glu286, Ala317, Asn330, Gln455, Arg459, Arg462, and Pro464 maintain the contacts more than 20 % of simulation time.

### 3.3. Molecular Docking of curcumin at RNA binding site of NS3 helicase

Next, we use a 5GJB crystal structure of NS3 helicase (Figure 3A) to dock the curcumin on RNA binding site. Similar to NTPase site, a grid was generated at the centroid of bound RNA at RNA binding site. Later, the Curcumin was docked at RNA binding site using XP docking approach. The docking result showed a docking score of −5.664 kcal/mol, and curcumin stabilizes itself at this site via four H-bonds through Arg388, Thr409, Asp540, and Ser601 residues (Figure 3B). One hydroxyl group of Curcumin forms two H-bonds with Arg388 and Thr409. The second hydroxyl group forms one hydrogen bond with Asp540. The fourth H-bond was formed between the keto group oxygen of curcumin and residue Ser601. Besides H-bond, the Curcumin stabilizes by a few hydrophobic interactions such as Pro292, Val366, Ile411, Pro432, Leu442, Val543, Pro542, and Leu541 (Figure 3C).

**Figure 3:**
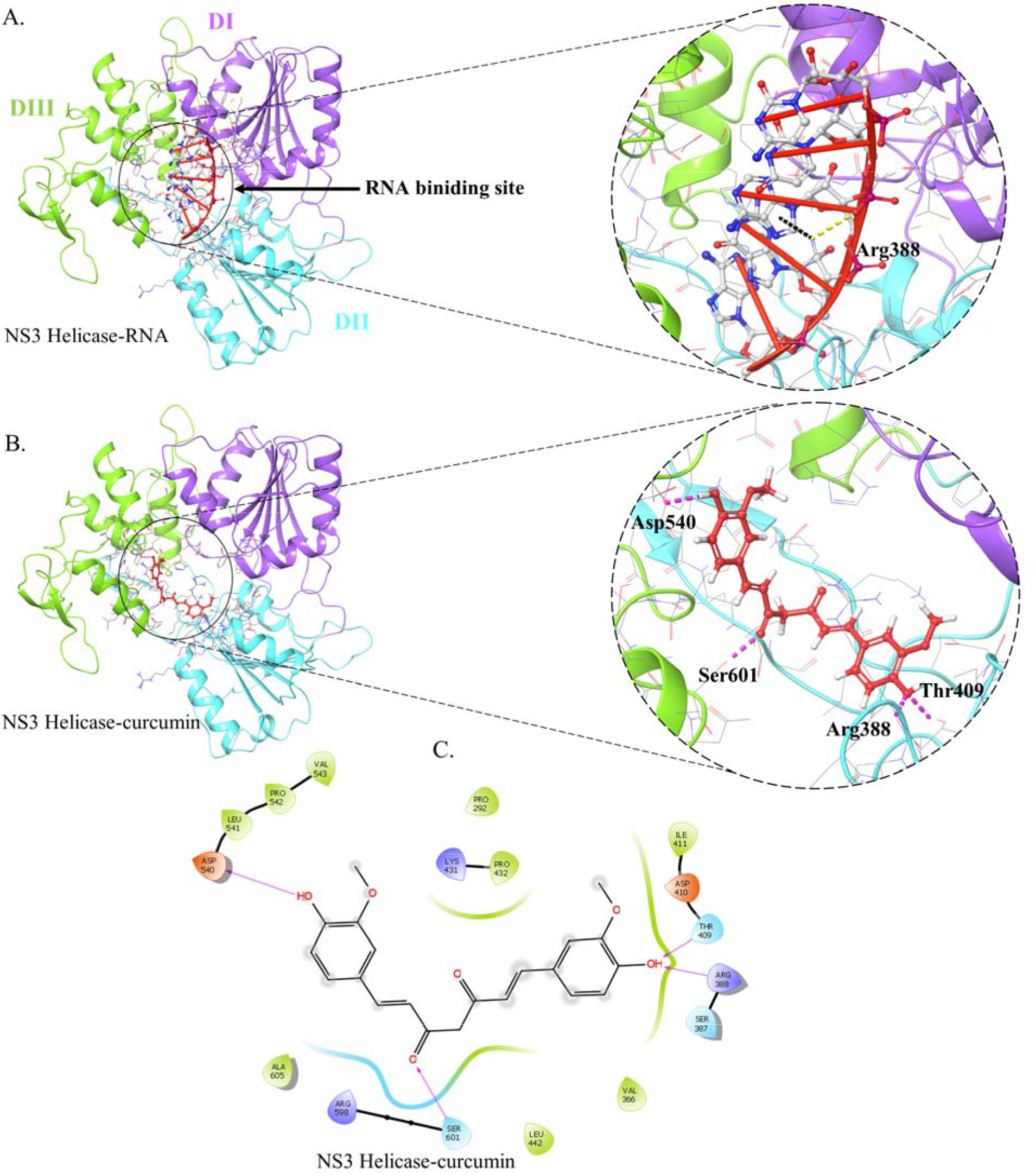
Molecular interaction of curcumin with RNA binding site of NS3 helicase in docking. (A) 5GJB crystal structure of NS3 helicase representing RNA binding site where bound RNA is shown in red colored. Three domains, DI (178-332), DII (333-481), and DII (482-617) of NS3 helicase, are represented in magenta, cyan, and green color, respectively. RNA stabilize at the RNA binding site by forming a salt bridge (yellow dashed line) and pi-cation bond (black dashed line) with Arg388 residue. (B) Three-dimensional pose of curcumin at the RNA binding site of NS3 helicase (5GJB) in docking. The pink dashed line represents the interaction of curcumin via H-bonds with Arg388, Thr409, Asp 540, and Ser601 of NS3 helicase. (C) Two-dimensional pose of curcumin at the RNA binding site of NS3 helicase representing its interaction with the amino acid residues. The pink arrow indicates the H-bond between the curcumin and NS3 helicase (5GJB) residues. The hydrophobic residues are represented in green color.

### 3.4. Curcumin forms stable interaction at RNA binding site of NS3 helicase

Further, we also checked the conformational flexibility and the stability of NS3 helicase (5GJB) upon binding curcumin to its RNA binding site. NS3 helicase with or without curcumin was subjected to MD simulation for 200 ns (Figure 4). During the simulation, C-α RMSD of NS3 helicase when curcumin was bound at its RNA binding site was observed to be 1.56-2.86 Å, a higher range than the unbound NS3 helicase (1.29-2.71 Å). The average RMSD of NS3 helicase is nearly similar, 2.16 Å or 2.17 Å in the absence or presence of curcumin, respectively (Figure 4A). These findings suggest that NS3 helicase exhibited a stable conformation when curcumin binds at the RNA binding site and did not show significant variation compared to unbound NS3 helicase. The ordered regions (helix or strand) of NS3 helicase showed lower RMSF than the disordered region (coil or loop), either in the presence or absence of curcumin (Figure 4B). Notably, when curcumin binds at the RNA binding site, the regions between 317-331 and 341-347 show low RMSF values. On the contrary, the region between residue 554-596 shows high RMSF values. Rg value of NS3 helicase bound with curcumin shows a slightly variable trend compared to unbound helicase (Figure 4C). The average Rg value of NS3 helicase is nearly similar in the presence (22.77 Å) or absence of curcumin (22.84 Å).

**Figure 4:**
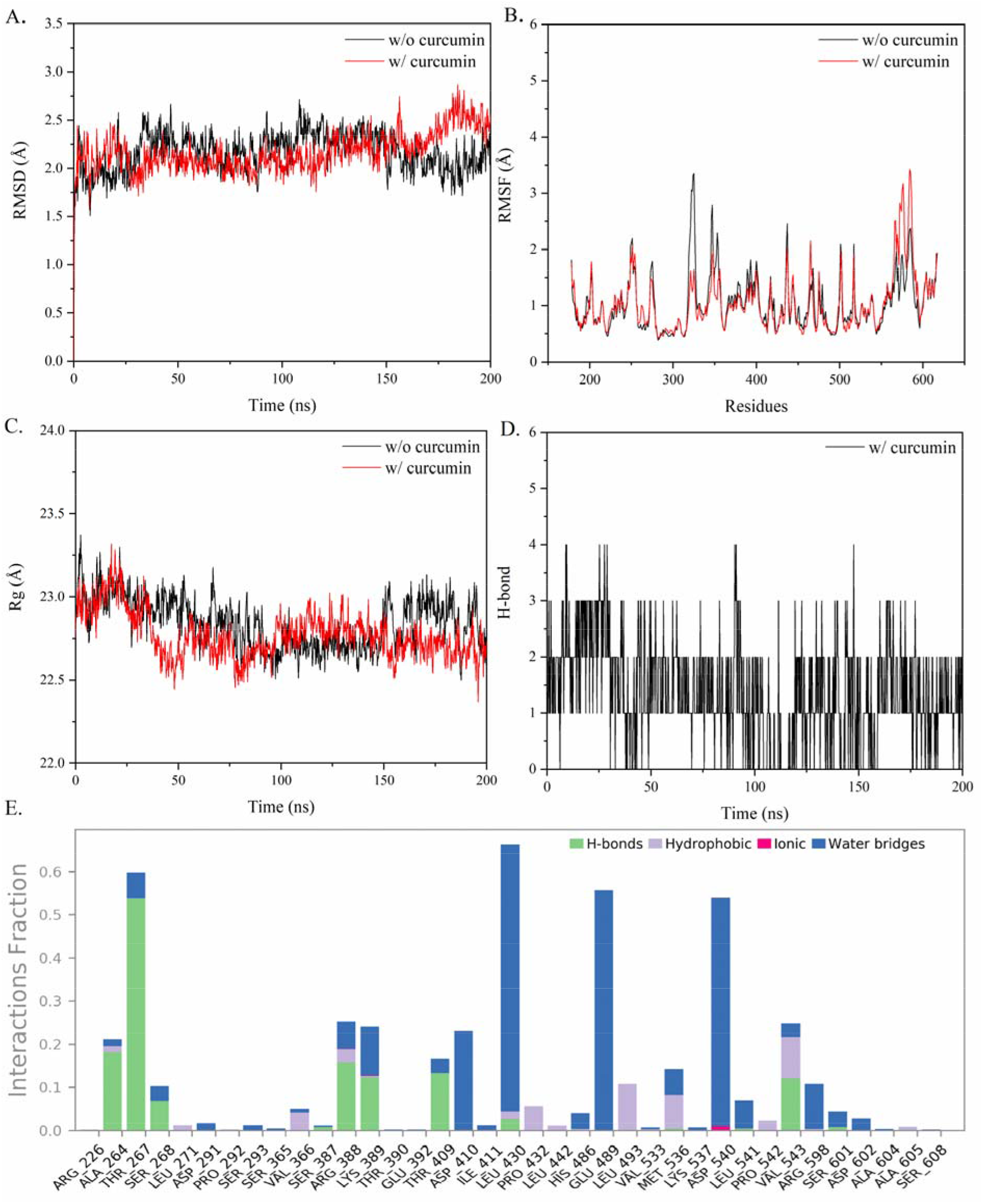
MD simulation of NS3 helicase when curcumin interacted with RNA-binding site. (A) The RMSD of NS3 helicase in the presence of curcumin (red colored line) and the absence of curcumin (black colored line). (B) The RMSF of NS3 helicase residue in the presence (red colored line) and the absence (black colored line) of curcumin. (C) The Rg of NS3 helicase in the presence (red colored line) and the absence (black colored line) of the curcumin. (D) The number of H-bond between curcumin and the RNA binding site of NS3 helicase throughout the simulation. (E) Fraction of interaction curcumin at the RNA binding site of NS3 helicase residues by H-bond, hydrophobic interaction, ionic bond, and water bridges during simulation. A fraction of 1 represents that the particular residue of helicase contacted with curcumin for 100 % of the simulation time. [5GJB crystal structure was used to dock the curcumin at the RNA binding site of NS3 helicase]

NS3 helicase has a 40.18 % total secondary structure element where the helix and strand are 25.59 % and 14.59 % during simulation (Figure S4). When bound with curcumin, there is little change in the secondary structure element (Figure S5) of NS3 helicase (Total SSE 38.81 %; Helix 25.95 % and Strand 12.86 %). Curcumin forms a maximum of four H-bonds in a few trajectory frames (Figure 4D). Curcumin shows contacts with 37 residues during the simulation and maintains the contacts with variable fractions of 200 ns simulation time (Figures 4E & S6). The residues Ala264, Thr267, Ser268, Ser387, Arg388, Lys389, Thr409, Leu430, Met536, Leu541, Val543, and Ser601 are involved in forming H-bond contacts with the curcumin. Arg388, Lys389, and Asp540 are involved in forming ionic bonds with curcumin. Arg226, Ala264, Leu271, Val366, Arg388, Lys389, Leu430, Pro430, Leu442, His486, Leu493, Val533, Met536, Pro542, Val543, Arg598, and Ala605 are involved in forming hydrophobic bonds with the curcumin. Ala264, Thr267, Ser268, Asp291, Ser293, Ser365, Val366, Ser387, Arg388, Lys389, Thr390, Glu392, Thr409, Asp410, Ile411, Leu430, His486, Glu489, Val533, Met536, Lys537, Asp540, Leu541, Val543, Arg598, Ser601, Asp602, Ala604, Ala605, and Ser608 are involved in forming water bridges with the curcumin. Among all the contacts between curcumin and NS3 helicase, Thr267, Leu430, Glu489, and Asp540 maintain the contacts for more than 50-70 % of the simulation time. Ala264, Ser268, Arg388, Lys389, Thr409, Asp410, Leu493, Met536, Val543, and Arg598 maintain contact between 10-30 % of the simulation time.

### 3.5. Curcumin inhibits NS3 helicase activity

In our computational studies, curcumin forms stable binding at NTPase site. Furthermore, we tested the potential of Curcumin to inhibit NTPase activity of NS3 helicase in vitro. First, we checked the NTPase activity of NS3 helicase protein in the absence and presence of variable curcumin concentrations (0, 25, 50, and 100 μM) with different substrate (ATP) concentrations (0-2500 μM) at 25°C in Tris buffer at pH 7.4 (Figure 5A). Then, data points were fitted with the Michaelis-Menten equation to estimate kinetic parameters (see Figure 5A and Table 2). The presence of curcumin decreases k_cat_ from 1.025 s^-1^ (no curcumin) to 0.202 s^-1^ (μM 100 curcumin), suggesting that the curcumin inhibits the helicase activity. Second, we tested NTPase activity in the presence of variable curcumin concentrations (0, 3.13, 6.25, 12.5, 25, 50, 100, 200 & 400 μM) at constant ATP concentration (1000 μM) to estimate IC_50_ for curcumin, which is 64.42 μM (Figure 5B).

**Figure 5:**
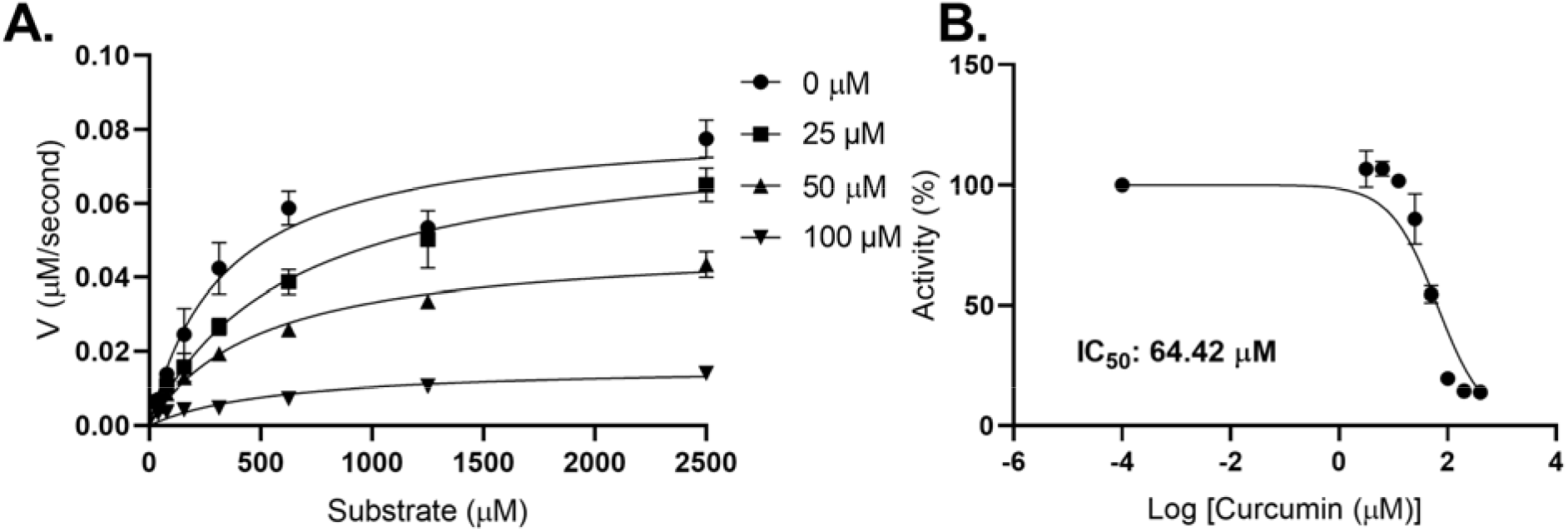
Inhibition of NS3 helicase NTPase activity by curcumin. (A) Plot of NS3 helicase NTPase activity in the presence of 0, 25, 50, and 100 _μ_M curcumin at variable substrate concentration, ATP (0-2500 _μ_M). The data point was fitted using the Michaelis-Menten equation (see Table 2 for kinetic parameters), and the error bar represents the standard error for triplicates. (B) Plot of NS3 helicase NTPase activity in the presence of a variable concentration of curcumin (0-400 _μ_M) with a constant ATP concentration (1000 _μ_M). The curcumin concentration at the X-axis was transformed into a logarithmic scale, and the data point was fitted using a dose-response inhibition equation. The error bar represents the standard error for triplicates.

**Table 1:** A summary of molecular docking result of curcumin.

| Protein (PDB ID) | Binding site | docking score (kcal/mol) | MMGBSA dG Bind (kcal/mol) | H-bond | π-π bond | π-cation bond | hydrophobic residues interaction |
| --- | --- | --- | --- | --- | --- | --- | --- |
| NS3 helicase (5GJC) | NTPase site | -4.93 | -18.94 | Phe418 | - | - | Pro196, Ala198, Ala317, Phe418, Ala420, Ile460, Pro464 |
| NS3 helicase (5GJB) | RNA binding site | -5.66 | -49.43 | Arg388, Thr409, Asp540, Ser601 | - | - | Pro292, Val366, Ile411, Pro432, Leu442, Leu541, Pro542, val543, Ala605 |
| NS2B-NS3 protease (5LC0) | Catalytic site | -4.17 | -30.95 | Tyr130 | Tyr161 | - | Phe84*, Trp50, Val72, Tyr150, Tyr161, Tyr130, Pro131, Ala132 |
| MTase (5KQR) | Druggable site as used in Sharma et al., 2020. [21] | -6.27 | -56.69 | Ser56, Arg84, Gly86, Glu111, Asp131, Val132 | - | - | Cys82, Trp87, Val130, Val132, Phe133, Ile147 |
| RdRp (5WZ3) | Druggable site as used in Sharma et al., 2020. [21] | -5.89 | -50.31 | Gln605, Tyr609, Arg794, Trp805 | - | Arg483 | Val606, Val607, Tyr609, Trp797, Ile799, Trp805, Met806 |

[\* represents the residue of NS2B in NS2B-NS3 protease]

**Table 2:** A summary of enzyme kinetic parameters of NS3 helicase in the presence of curcumin.

| Kinetic parameter | DMSO | Curcumin |  |  |
| --- | --- | --- | --- | --- |
| | 1% | 25 μM | 50 μM | 100 μM |
| $V_{\max}$ (μM/second) | $0.082 \pm 0.005$ | $0.079 \pm 0.004$ | $0.049 \pm 0.002$ | $0.016 \pm 0.0017$ |
| $K_m$ (μM) | $336.3 \pm 61.97$ | $633.9 \pm 84.81$ | $497.1 \pm 55.36$ | $564.7 \pm 158.1$ |
| $k_{\text{cat}}$ ($\text{second}^{-1}$) | $1.025 \pm 0.06$ | $0.992 \pm 0.052$ | $0.62 \pm 0.025$ | $0.202 \pm 0.02$ |
| $R^2$ | 0.93 | 0.97 | 0.97 | 0.82 |

### 3.6. Curcumin also interacts with the druggable site of ZIKV MTase & RdRp

We have also checked the interaction of curcumin with other ZIKV enzymes like NS2B-NS3 protease, MTase, and RdRp (Figure 6). We used 5LC0, 5KQR, and 5WZ3 crystal structures for NS2B-NS3 protease (Figure 6A), MTase (Figure 6B), and RdRp (Figure 6C), respectively. Curcumin interacts at the protease active site via π-π contact with Tyr161 and H-bond with Tyr130 of NS3 protease. NS2B-NS3 protease-curcumin complex shows a docking score of −4.17 kcal/mol. In the case of MTase, curcumin at its druggable site interacts via seven H-bonds with Ser56, Arg84, Gly86, Glu111, Val132, and Asp131. MTase-curcumin complex shows docking score of −6.27 kcal/mol. Curcumin shows multiple interactions at the druggable site of RdRp via five H-bonds through Gln605, Tyr609, Arg794, and Trp805. RdRp-curcumin complex shows docking score of −5.89 kcal/mol. Here, docking of the curcumin suggests a good binding affinity towards the druggable site of these enzymes. However, further experimental validation needs to be done by NMR and the crystal structure of enzyme-curcumin complex.

**Figure 6:**
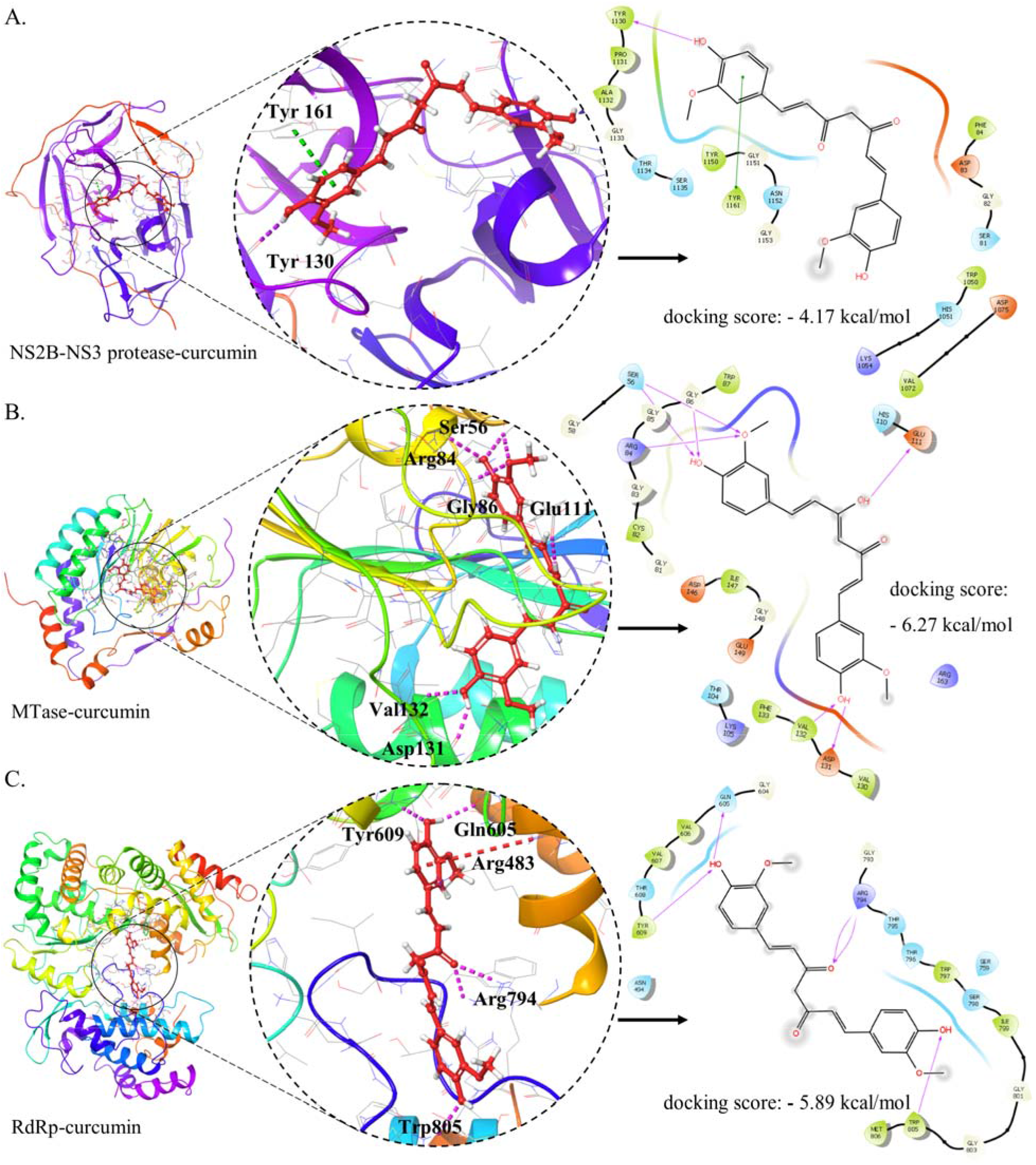
Molecular interaction of curcumin at NS2B-NS3 protease, MTase, and RdRp protein of Zika virus in docking. (A) Interaction of curcumin at NS2B-NS3 protease (5LC0) active site. The blue ribbon in 3D pose represents NS3 protease part, and the red ribbon represents NS2B (residues 49-95). For clarity, NS2B is 49-95, and NS3 is 1001 to 1170, corresponding to 1 to 170. (B) Molecular interaction of curcumin at the MTase (5KQR). (C) Molecular interaction of curcumin at the RdRp (5WZ3). Curcumin is represented in red in all the panels. A pink dashed line and arrow represent the H-bond, a green dashed line represents a π-π bond, and a red dashed line represents a π-cation bond.

## 4. DISCUSSION

To meet the demands for treatment and prepare for the ever-present threat of future large-scale epidemics, research efforts must be sustained in drug discovery so that improved strategies will become available for treating and preventing ZIKV infections. Various compounds with potentially beneficial properties, including natural products, nucleoside analogs, peptides, small molecules, antibiotics, and others, have been identified previously as possible inhibitors of ZIKV in both *in vitro* and *in vivo* or only *in vitro* studies [24]. These inhibitors have demonstrated the ability to inhibit entry of the virus by targeting the envelope proteins or inhibit the replication or assembly of the virus by targeting the non-structural proteins that include protease, helicase, and polymerase [24]. For an inhibitor discovery, identifying a specific target plays a vital role in the success of a lead molecule, and further understanding the molecular interaction of these molecules at their target will be very useful in identifying novel inhibitor molecules against the Zika virus.

Previously, natural products have been used in drug discovery research for infectious diseases, cardiovascular diseases, cancer, and sclerosis [25]. In this regard, some natural products and their analogs, like nanchangmycin, gossypol, conessine, digitonin, baicalein, baicalin, emodin, berberine, theaflavin, EGCG, curcumin, have been identified as potential inhibitor candidates for ZIKV infection [24]. It has been noted that most of these inhibitors target the ZIKV entry step to the host cells to inhibit ZIKV infection. Moreover, baicalein, baicalin, and Theaflavin can also target ZIKV NS5 protein. EGCG can target NS2B-NS3 protease [22] and NS3 helicase [11] of ZIKV in addition to the viral entry [20,26]. Similarly, the natural product curcumin potentially targets NS3 protease and inhibits its functional activity [27]. Notably, its antiviral properties and targeting of NS3 protease activity have also been discovered in DENV [28,29]. Looking at the previous literature, we find that curcumin inhibits the replication of a wide range of viruses [16]. Further, in this study, we explored the molecular interaction of curcumin with ZIKV NS3 helicase and its potential to inhibit NS3 helicase activity.

For ZIKV, NS3 helicase is essential for genome replication, where NTPase site acts as a binding site for ATP, which provides energy for RNA unwinding at this RNA binding site [2]. Therefore, blocking these two sites will lead to the inhibition of NS3 helicase activity and thus may inhibit ZIKV replication. Our study finds that curcumin forms stable interactions at both ATP and RNA binding sites of NS3 helicase by docking and simulation, suggesting that curcumin can block the functional activity of NS3 helicase. So, further, we assessed the inhibition potential of curcumin by analyzing NTPase activity of NS3 helicase. We found that curcumin inhibits the functional activity of NS3 helicase with inhibition concentration in the micromolar range. In line with this finding, our group previously identified the NS3 helicase as a target for EGCG, a green tea molecule, which reduces the NS3 helicase activity in the nanomolar range [11]. Therefore, the mechanistic insight into the action of curcumin will enlighten further drug discovery research to identify similar molecules that can target the NS3 helicase of the Zika virus and act as a broad-spectrum inhibitor for other viruses of the same family.

## Author contribution

**Ankur Kumar:** Conceptualization, Methodology, Investigation, Analysis, Writing-Original drafts. **Aparna Bhardwaj:** Investigation. **Richa Joshi:** Investigation. **Taniya Bhardwaj:** Writing-Original drafts. **Rajanish Giri:** Conceptualization, Supervision, Writing-Review & Editing, Funding acquisition.

## Acknowledgments

We want to thank IIT Mandi for providing research space and facilities. This work is supported by Indian Council of Medical Research (52/04/2020/BIO/BMS) grant to RG. RG is also grateful to Science and Engineering Research Board (SERB) grant, India (CRG/2019/005603); IYBA Award (BT/11/IYBA/2018/06) grant, Department of Biotechnology (DBT), India; Indian Council of Medical Research (58/6/2020/PHA/BMS), and MHRD-SPARC (SPARC/2018-2019/P37/SL).

## References

[1] M.Y.F. Tay, W.G. Saw, Y. Zhao, K.W.K. Chan, D. Singh, Y. Chong, J.K. Forwood, E.E. Ooi, G. Grüber, J. Lescar, D. Luo, S.G. Vasudevan, The C-terminal 50 Amino Acid Residues of Dengue NS3 Protein Are Important for NS3-NS5 Interaction and Viral Replication *, J. Biol. Chem. 290 (2015) 2379–2394. 10.1074/JBC.M114.607341.

[2] D. Luo, S.G. Vasudevan, J. Lescar, The flavivirus NS2B–NS3 protease–helicase as a target for antiviral drug development, Antiviral Res. 118 (2015) 148–158. 10.1016/J.ANTIVIRAL.2015.03.014.

[3] H. Tian, X. Ji, X. Yang, W. Xie, K. Yang, C. Chen, C. Wu, H. Chi, Z. Mu, Z. Wang, H. Yang, The crystal structure of Zika virus helicase: basis for antiviral drug design, Protein Cell. 7 (2016) 450–454. 10.1007/S13238-016-0275-4.

[4] H. Tian, X. Ji, X. Yang, Z. Zhang, Z. Lu, K. Yang, C. Chen, Q. Zhao, H. Chi, Z. Mu, W. Xie, Z. Wang, H. Lou, H. Yang, Z. Rao, Structural basis of Zika virus helicase in recognizing its substrates, Protein Cell. 7 (2016) 562–570. 10.1007/S13238-016-0293-2/FIGURES/5.

[5] D. Luo, T. Xu, R.P. Watson, D. Scherer-Becker, A. Sampath, W. Jahnke, S.S. Yeong, C.H. Wang, S.P. Lim, A. Strongin, S.G. Vasudevan, J. Lescar, Insights into RNA unwinding and ATP hydrolysis by the flavivirus NS3 protein, EMBO J. 27 (2008) 3209– 3219. 10.1038/EMBOJ.2008.232.

[6] C.A. Belon, D.N. Frick, Helicase inhibitors as specifically targeted antiviral therapy for hepatitis C, 10.2217/Fvl.09.7. 4 (2009) 277–293. 10.2217/FVL.09.7.

[7] G. Maga, S. Gemma, C. Fattorusso, G.A. Locatelli, S. Butini, M. Persico, G. Kukreja, M.P. Romano, L. Chiasserini, L. Savini, E. Novellino, V. Nacci, S. Spadari, G. Campiani, Specific targeting of hepatitis C virus NS3 RNA helicase. Discovery of the potent and selective competitive nucleotide-mimicking inhibitor QU663, Biochemistry. 44 (2005) 9637–9644. 10.1021/BI047437U/SUPPL_FILE/BI047437USI20050404_085959.PDF.

[8] I. Briguglio, S. Piras, P. Corona, A. Carta, Inhibition of RNA Helicases of ssRNA + Virus Belonging to Flaviviridae, Coronaviridae and Picornaviridae Families, Int. J. Med. Chem. 2011 (2011) 1–22. 10.1155/2011/213135.

[9] N.L. Sweeney, A.M. Hanson, S. Mukherjee, J. Ndjomou, B.J. Geiss, J.J. Steel, K.J. Frankowski, K. Li, F.J. Schoenen, D.N. Frick, Benzothiazole and Pyrrolone Flavivirus Inhibitors Targeting the Viral Helicase, ACS Infect. Dis. 1 (2015) 140–148. 10.1021/ID5000458/SUPPL_FILE/ID5000458_SI_001.PDF.

[10] E. Mastrangelo, M. Pezzullo, T. De burghgraeve, S. Kaptein, B. Pastorino, K. Dallmeier, X. De lamballerie, J. Neyts, A.M. Hanson, D.N. Frick, M. Bolognesi, M. Milani, Ivermectin is a potent inhibitor of flavivirus replication specifically targeting NS3 helicase activity: new prospects for an old drug, J. Antimicrob. Chemother. 67 (2012) 1884. 10.1093/JAC/DKS147.

[11] D. Kumar, N. Sharma, M. Aarthy, S.K. Singh, R. Giri, Mechanistic Insights into Zika Virus NS3 Helicase Inhibition by Epigallocatechin-3-Gallate, ACS Omega. 5 (2020) 11217–11226. 10.1021/ACSOMEGA.0C01353/ASSET/IMAGES/MEDIUM/AO0C01353_M001.GIF.

[12] S. Zorofchian Moghadamtousi, H. Abdul Kadir, P. Hassandarvish, H. Tajik, S. Abubakar, K. Zandi, A review on antibacterial, antiviral, and antifungal activity of curcumin, Biomed Res. Int. 2014 (2014). 10.1155/2014/186864.

[13] K.R. Kahkhaie, A. Mirhosseini, A. Aliabadi, A. Mohammadi, M.J. Mousavi, S.M. Haftcheshmeh, T. Sathyapalan, A. Sahebkar, Curcumin: a modulator of inflammatory signaling pathways in the immune system, Inflammopharmacology 2019 275. 27 (2019) 885–900. 10.1007/S10787-019-00607-3.

[14] S.H. Mun, D.K. Joung, Y.S. Kim, O.H. Kang, S.B. Kim, Y.S. Seo, Y.C. Kim, D.S. Lee, D.W. Shin, K.T. Kweon, D.Y. Kwon, Synergistic antibacterial effect of curcumin against methicillin-resistant Staphylococcus aureus, Phytomedicine. 20 (2013) 714–718. 10.1016/J.PHYMED.2013.02.006.

[15] A. Zia, T. Farkhondeh, A.M. Pourbagher-Shahri, S. Samarghandian, The role of curcumin in aging and senescence: Molecular mechanisms, Biomed. Pharmacother. 134 (2021) 111119. 10.1016/J.BIOPHA.2020.111119.

[16] M.R. Jennings, R.J. Parks, Curcumin as an Antiviral Agent, Viruses. 12 (2020). 10.3390/V12111242.

[17] A. Balasubramanian, R. Pilankatta, T. Teramoto, A.M. Sajith, E. Nwulia, A. Kulkarni, R. Padmanabhan, Inhibition of dengue virus by curcuminoids, Antiviral Res. 162 (2019) 71–78. 10.1016/J.ANTIVIRAL.2018.12.002.

[18] H. Tian, X. Ji, X. Yang, Z. Zhang, Z. Lu, K. Yang, C. Chen, Q. Zhao, H. Chi, Z. Mu, W. Xie, Z. Wang, H. Lou, H. Yang, Z. Rao, Structural basis of Zika virus helicase in recognizing its substrates, Protein Cell. 7 (2016) 562. 10.1007/S13238-016-0293-2.

[19] A. Kumar, D. Kumar, P. Kumar, B.L. Jones, I.U. Mysorekar, R. Giri, Discovery And Characterization of Small Molecule Inhibitors of Zika Virus Replication, BioRxiv. (2022) 2022.12.15.520558. 10.1101/2022.12.15.520558.

[20] N. Sharma, A. Murali, S.K. Singh, R. Giri, Epigallocatechin gallate, an active green tea compound inhibits the Zika virus entry into host cells via binding the envelope protein, Int. J. Biol. Macromol. 104 (2017) 1046–1054. 10.1016/J.IJBIOMAC.2017.06.105.

[21] N. Sharma, O. Prosser, P. Kumar, A. Tuplin, R. Giri, Small molecule inhibitors possibly targeting the rearrangement of Zika virus envelope protein, Antiviral Res. 182 (2020) 104876. 10.1016/J.ANTIVIRAL.2020.104876.

[22] R. Yadav, C. Selvaraj, M. Aarthy, P. Kumar, A. Kumar, S.K. Singh, R. Giri, Investigating into the molecular interactions of flavonoids targeting NS2B-NS3 protease from ZIKA virus through *in-silico* approaches, J. Biomol. Struct. Dyn. (2020) 1–13. 10.1080/07391102.2019.1709546.

[23] S. Genheden, U. Ryde, The MM/PBSA and MM/GBSA methods to estimate ligand-binding affinities, Expert Opin. Drug Discov. 10 (2015) 449. 10.1517/17460441.2015.1032936.

[24] A. Kumar, D. Kumar, J. Jose, R. Giri, I.U. Mysorekar, Drugs to limit Zika virus infection and implication for maternal-fetal health, Front. Virol. 0 (2022) 64. 10.3389/FVIRO.2022.928599.

[25] A.G. Atanasov, S.B. Zotchev, V.M. Dirsch, I.E. Orhan, M. Banach, J.M. Rollinger, D. Barreca, W. Weckwerth, R. Bauer, E.A. Bayer, M. Majeed, A. Bishayee, V. Bochkov, G.K. Bonn, N. Braidy, F. Bucar, A. Cifuentes, G. D’Onofrio, M. Bodkin, M. Diederich, A.T. Dinkova-Kostova, T. Efferth, K. El Bairi, N. Arkells, T.P. Fan, B.L. Fiebich, M. Freissmuth, M.I. Georgiev, S. Gibbons, K.M. Godfrey, C.W. Gruber, J. Heer, L.A. Huber, E. Ibanez, A. Kijjoa, A.K. Kiss, A. Lu, F.A. Macias, M.J.S. Miller, A. Mocan, R. Müller, F. Nicoletti, G. Perry, V. Pittalà, L. Rastrelli, M. Ristow, G.L. Russo, A.S. Silva, D. Schuster, H. Sheridan, K. Skalicka-Woźniak, L. Skaltsounis, E. Sobarzo-Sánchez, D.S. Bredt, H. Stuppner, A. Sureda, N.T. Tzvetkov, R.A. Vacca, B.B. Aggarwal, M. Battino, F. Giampieri, M. Wink, J.L. Wolfender, J. Xiao, A.W.K. Yeung, G. Lizard, M.A. Popp, M. Heinrich, I. Berindan-Neagoe, M. Stadler, M. Daglia, R. Verpoorte, C.T. Supuran, Natural products in drug discovery: advances and opportunities, Nat. Rev. Drug Discov. 2021 203. 20 (2021) 200–216. 10.1038/s41573-020-00114-z.

[26] B.M. Carneiro, M.N. Batista, A.C.S. Braga, M.L. Nogueira, P. Rahal, The green tea molecule EGCG inhibits Zika virus entry, Virology. 496 (2016) 215–218. 10.1016/J.VIROL.2016.06.012.

[27] A. Roy, L. Lim, S. Srivastava, Y. Lu, J. Song, Solution conformations of Zika NS2B-NS3pro and its inhibition by natural products from edible plants, PLoS One. 12 (2017) e0180632. 10.1371/JOURNAL.PONE.0180632.

[28] L. Padilla-S, A. Rodríguez, M.M. Gonzales, J.C. Gallego-G, J.C. Castaño-O, Inhibitory effects of curcumin on dengue virus type 2-infected cells in vitro, Arch. Virol. 159 (2014) 573–579. 10.1007/S00705-013-1849-6/FIGURES/3.

[29] L. Lim, M. Dang, A. Roy, J. Kang, J. Song, Curcumin Allosterically Inhibits the Dengue NS2B-NS3 Protease by Disrupting Its Active Conformation, ACS Omega. 5 (2020) 25677–25686. 10.1021/ACSOMEGA.0C00039/ASSET/IMAGES/LARGE/AO0C00039_0007.JPEG.

