## Supplemental data for "Curcumin Inhibits Zika Virus NS3 Helicase Activity"

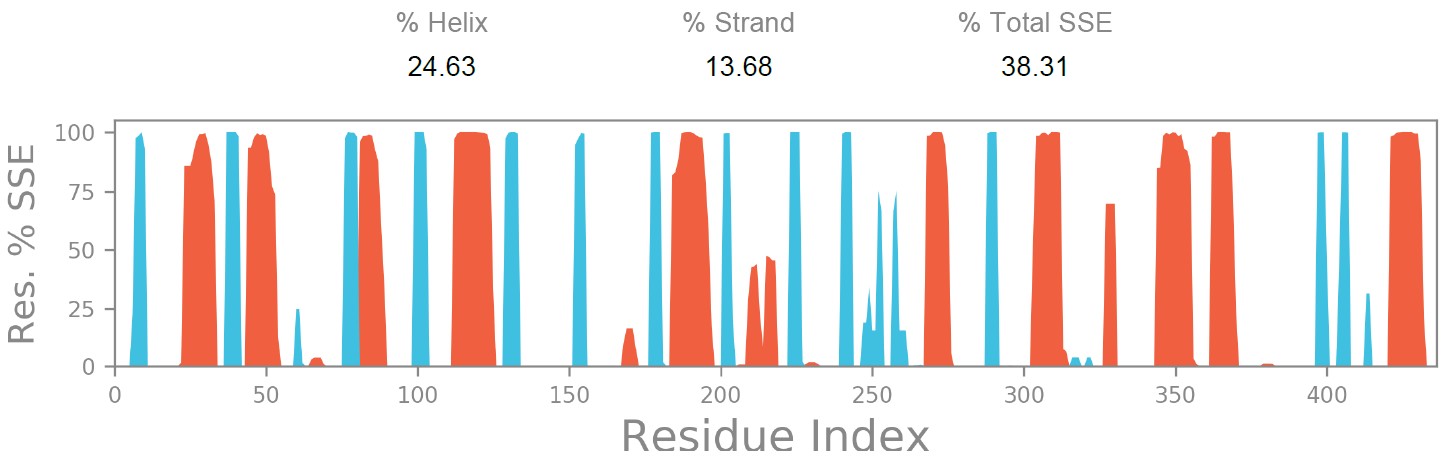

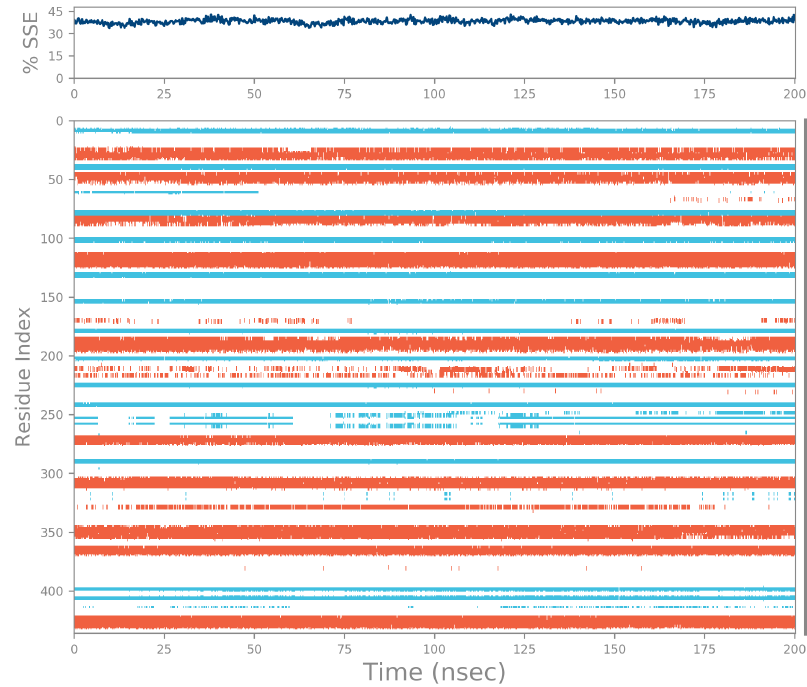

**Figure S1:** SSE distribution over residue index (upper panel), SSE composition of each trajectory frame (middle), and SSE assignment by each residue (bottom) of NS3 helicase (5GJC) in the absence of curcumin over the course of simulation of 200 ns. The red and blue colored segment in the plot represents the protein secondary structure element like helix and strand, respectively. “nsec” represents the time in nanoseconds.

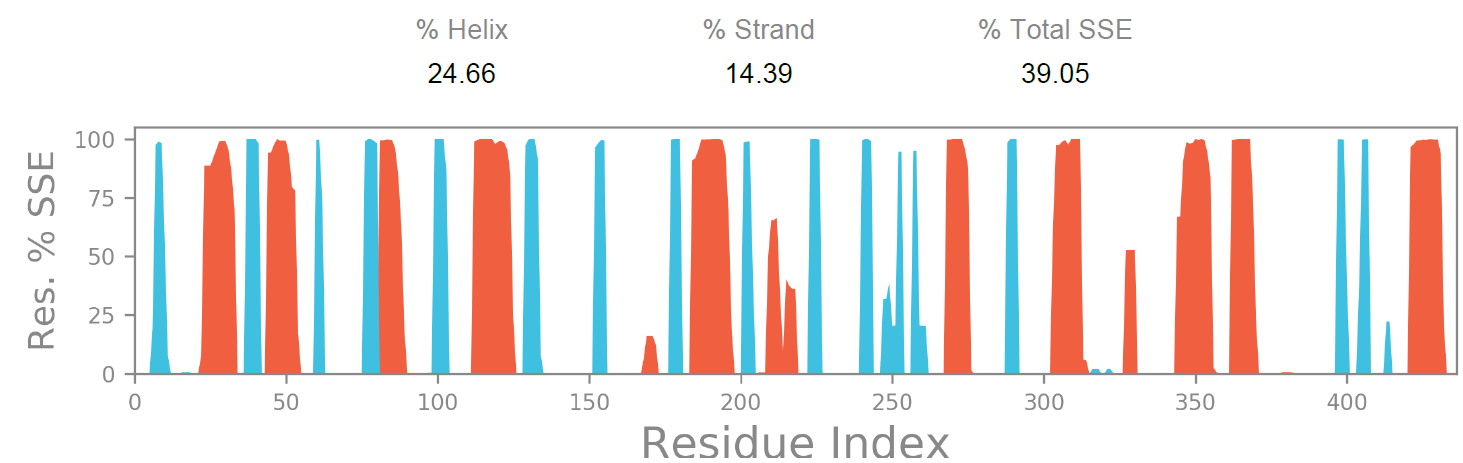

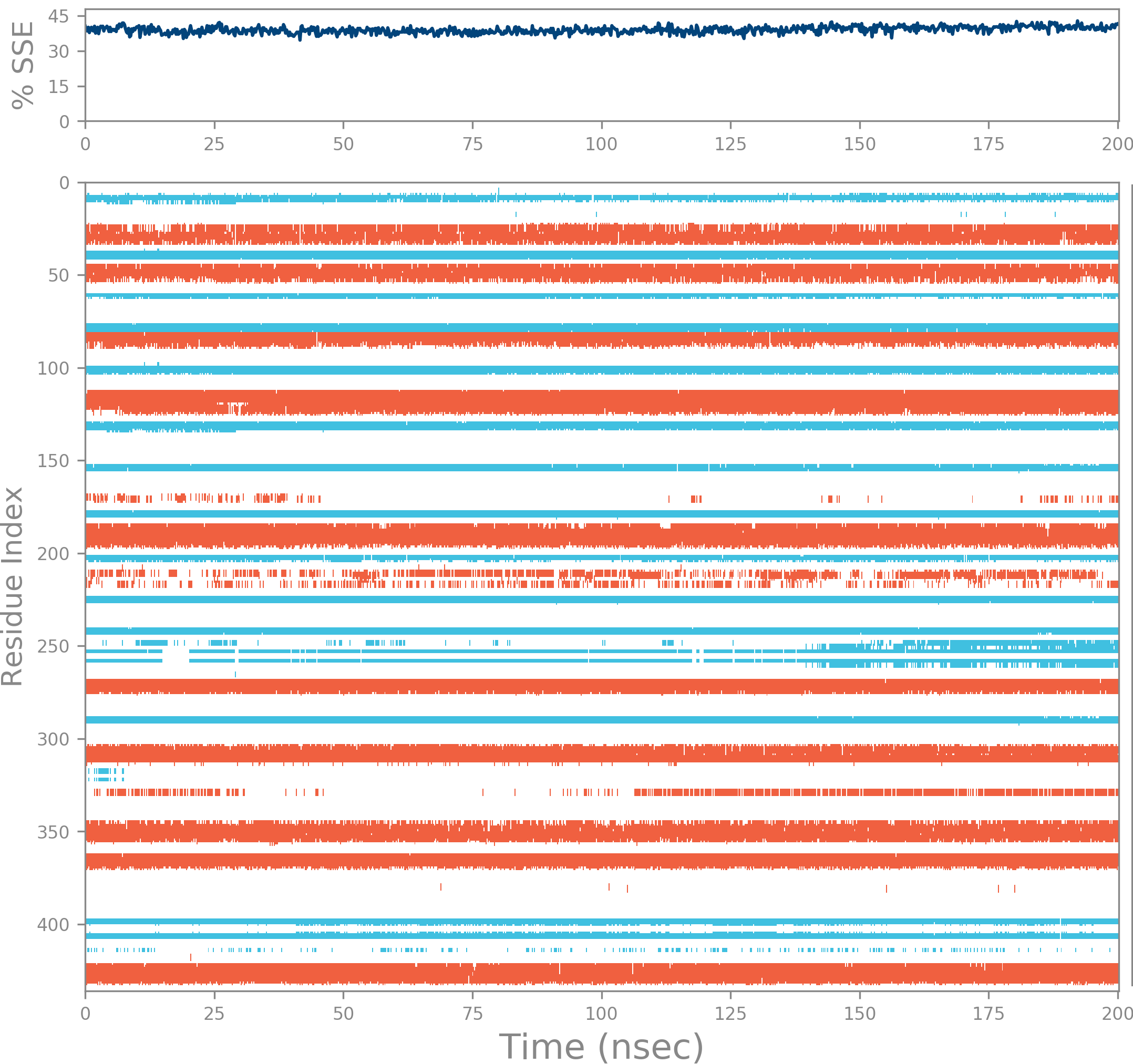

**Figure S2:** SSE distribution over residue index (upper panel), SSE composition of each trajectory frame (middle), and SSE assignment by each residue (bottom) of NS3 helicase (5GJC) in the presence of curcumin at NTPase site over the course of simulation of 200 ns. The red and blue colored segment in the plot represents the protein secondary structure element like helix and strand, respectively. “nsec” represents the time in nanoseconds.

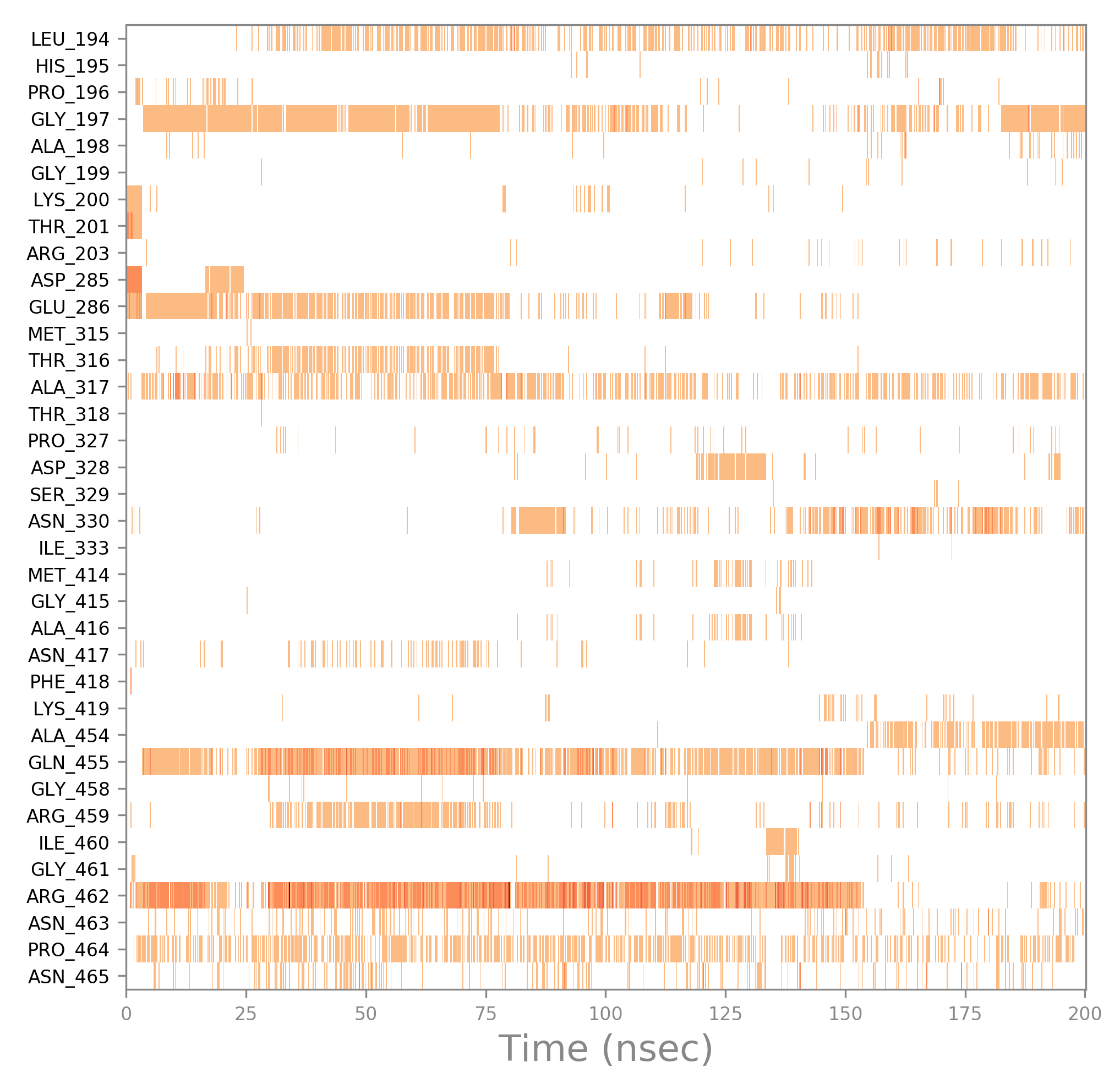
**Figure S3:** Molecular interaction of curcumin with the amino acid residues at the NTPase site of NS3 helicase (5GJC) over the course of simulation. The orange shade represents the interaction type (either H-bond or hydrophobic or ionic or water bridges). The dark orange color shade represents more than one type of interaction by that residue (either H-bond or hydrophobic or ionic or water bridges). “nsec” represents the time in nanoseconds.

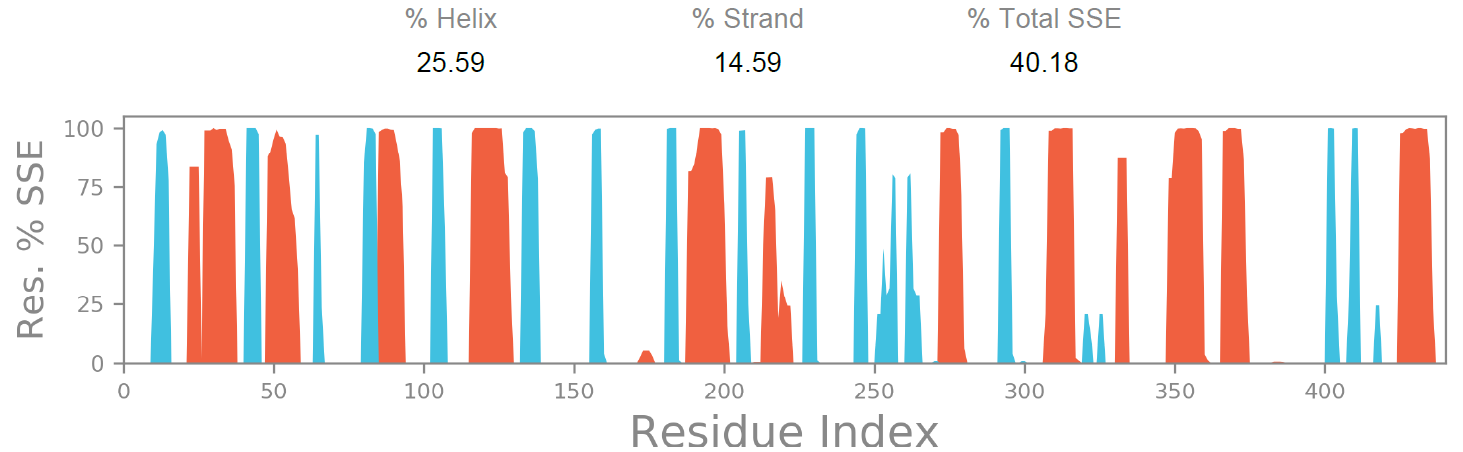

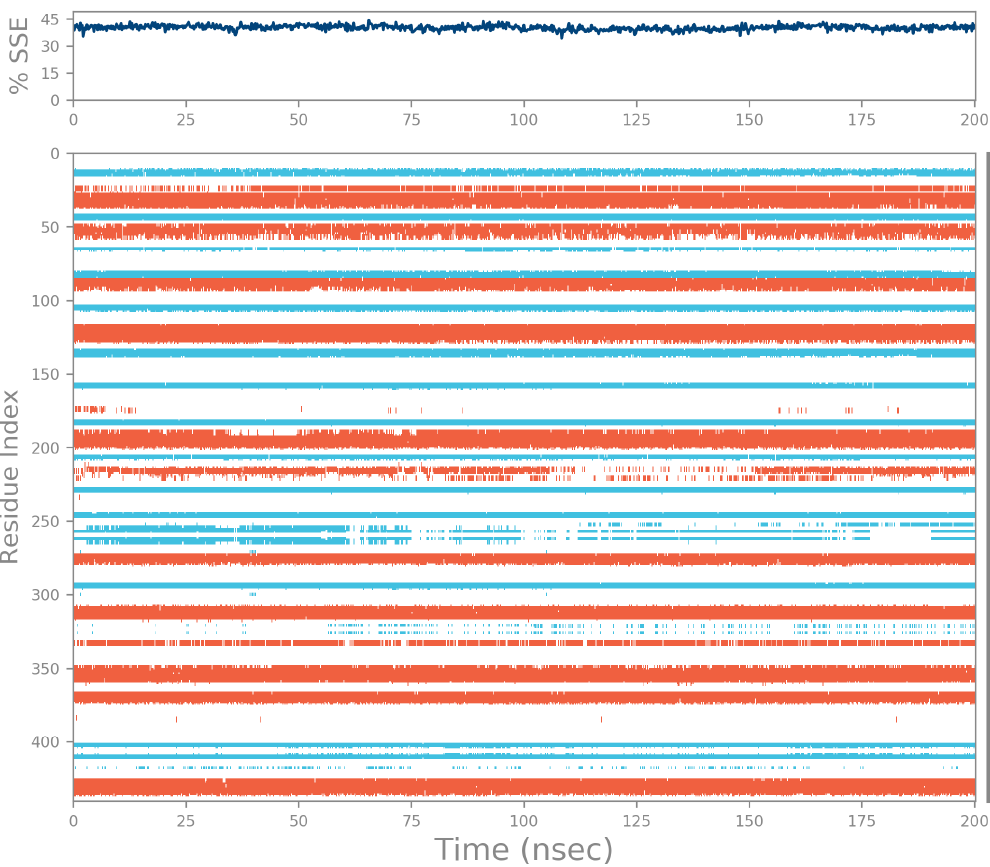

**Figure S4:** SSE distribution over residue index (upper panel), SSE composition of each trajectory frame (middle), and SSE assignment by each residue (bottom) of NS3 helicase (5GJB) in the absence of curcumin at RNA binding site over the course of simulation of 200 ns. The red and blue colored segment in the plot represents the protein secondary structure element like helix and strand, respectively. “nsec” represents the time in nanoseconds.

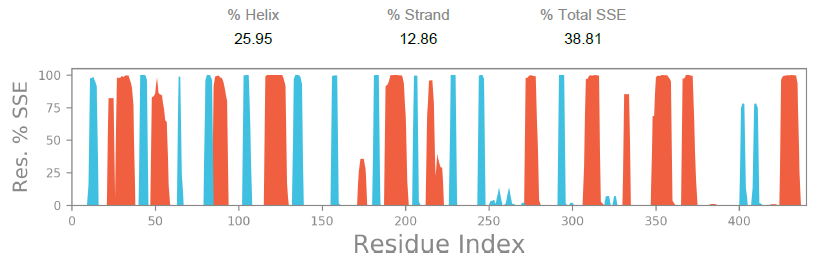

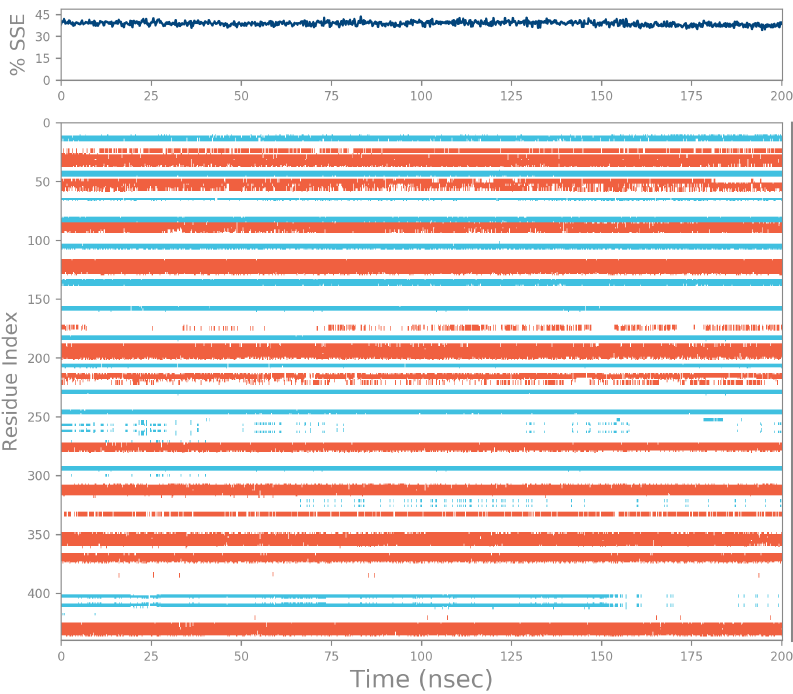

**Figure S5:** SSE distribution over residue index (upper panel), SSE composition of each trajectory frame (middle), and SSE assignment by each residue (bottom) of NS3 helicase (5GJB) in the presence of curcumin at RNA binding site over the course of simulation of 200 ns. The red and blue colored segment in the plot represents the protein secondary structure element like helix and strand, respectively. “nsec” represents the time in nanoseconds.

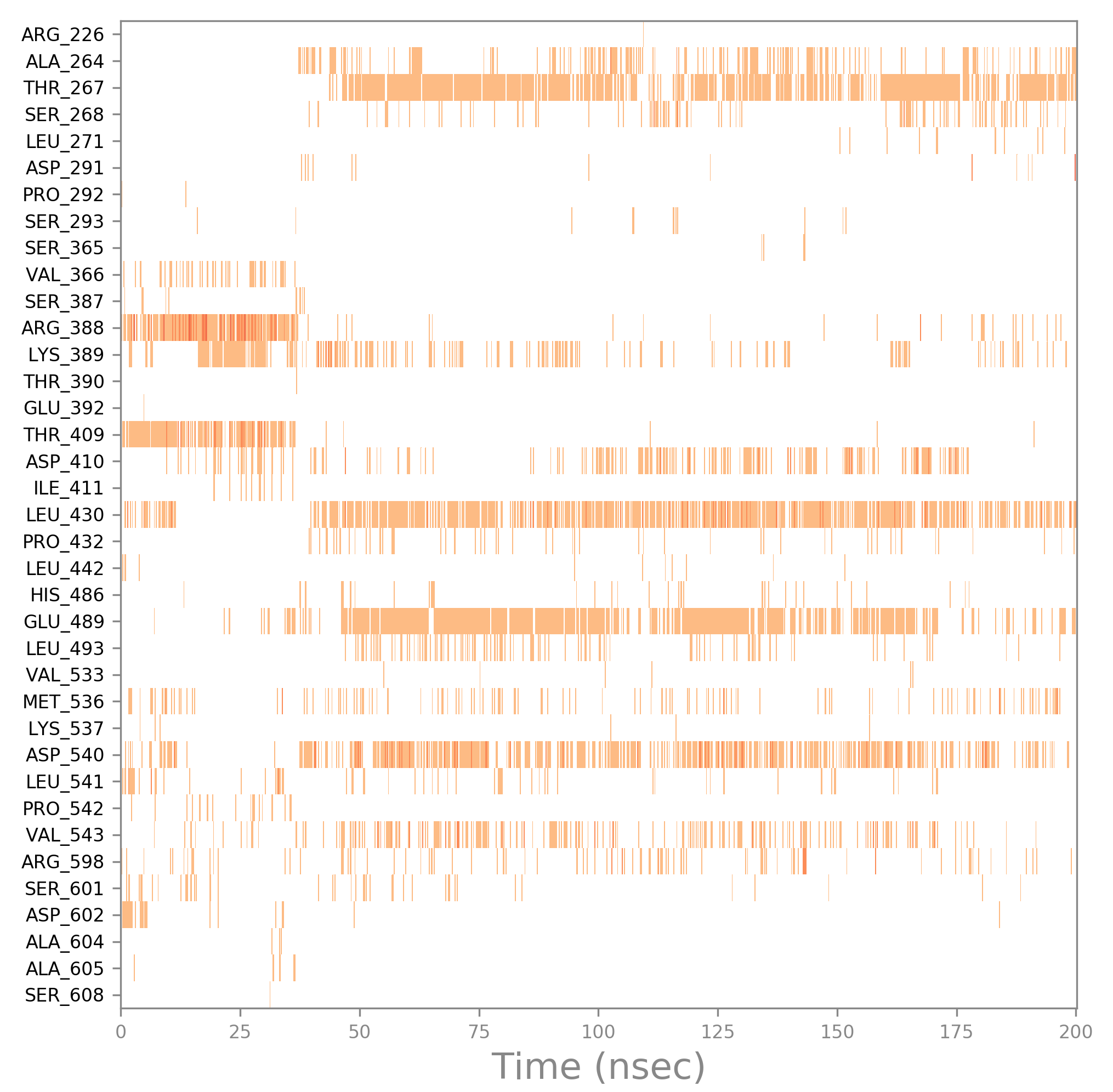

**Figure S6:** Molecular interaction of curcumin with the amino acid residues at the RNA binding site of NS3 helicase (5GJB) over the course of the simulation. The orange shade represents the interaction type (either H-bond or hydrophobic or ionic or water bridges). The dark orange color shade represents more than one type of interaction by that residue (either H-bond or hydrophobic or ionic or water bridges). “nsec” represents the time in nanoseconds.
